# A registration strategy to characterize DTI-observed changes in skeletal muscle architecture due to passive shortening

**DOI:** 10.1101/2024.04.11.589123

**Authors:** Melissa T. Hooijmans, Carly A. Lockard, Xingyu Zhou, Crystal Coolbaugh, Roberto Pineda Guzman, Mariana E. Kersh, Bruce M. Damon

## Abstract

Skeletal muscle architecture is a key determinant of muscle function. Architectural properties such as fascicle length, pennation angle, and curvature can be characterized using Diffusion Tensor Imaging (DTI), but acquiring these data during a contraction is not currently feasible. However, an image registration-based strategy may be able to convert muscle architectural properties observed at rest to their contracted state. As an initial step toward this long-term objective, the aim of this study was to determine if an image registration strategy could be used to convert the whole-muscle average architectural properties observed in the extended joint position to those of a flexed position, following passive rotation. DTI and high-resolution fat/water scans were acquired in the lower leg of seven healthy participants on a 3T MR system in +20° (plantarflexion) and −10° (dorsiflexion) foot positions. The diffusion and anatomical images from the two positions were used to propagate DTI fiber-tracts from seed points along a mesh representation of the aponeurosis of fiber insertion. The −10° and +20° anatomical images were registered and the displacement fields were used to transform the mesh and fiber-tracts from the +20° to the −10° position. Student’s paired *t*-tests were used to compare the mean architectural parameters between the original and transformed fiber-tracts. The whole-muscle average fiber-tract length, pennation angle, curvature, and physiological cross-sectional areas estimates did not differ significantly. DTI fiber-tracts in plantarflexion can be transformed to dorsiflexion position without significantly affecting the average architectural characteristics of the fiber-tracts. In the future, a similar approach could be used to evaluate muscle architecture in a contracted state.

## Introduction

The diversity of human motion, including movements that vary in velocity, degree of precision, force, and duration, requires a broad range of outputs from the musculoskeletal system. The mechanisms for these varied outputs include neural control strategies (the activation patterns of multiple muscles; the motor unit recruitment and rate coding strategies within a muscle) and the activated muscles’ metabolic, structural, and mechanical properties (1, 2). Several determinants of a muscle’s mechanical properties include its gross morphology; its internal structure (or architecture, as represented by the pennation angle, length, and curvature of its fibers); and the structural and mechanical properties of the collagen-rich structures (such as tendons and aponeuroses) with which it interacts (3). Skeletal muscle architecture can be non-invasively characterized using Diffusion Tensor Imaging (DTI), a magnetic resonance imaging (MRI) technique that measures the reduced and anisotropic diffusion of water molecules (4, 5). By measuring the anisotropic movement of water molecules, it is possible to derive information about the micro-structure and architecture of healthy and diseased muscle (6–23).

To date, DTI has primarily been applied in non-contracting, healthy muscle tissue. From such measurements, the microstructural and architectural properties of resting muscle have been characterized, including following passive joint motions (14, 16, 24). Furthermore, these microstructural and architectural properties have been used as inputs to musculoskeletal models to predict changes in muscle function in healthy, diseased, and injured muscle (25–30). DTI has intrinsic three-dimensional sensitivity, whole-muscle coverage, and the potential to be integrated with other forms of functionally relevant MRI contrast. With this approach, it may be possible to answer previously unrealizable questions in the study of muscle structure-function relationships.

To realize such possibilities, DTI data would ideally be acquired during a muscle contraction; but this is challenging due to long scan times required for muscle DTI, the potential for motion artefacts, and signal voids that may occur when diffusion-weighted images are acquired during contraction (31–33). This challenge is amplified by the fact that whole muscle DTI is required to provide a complete understanding of the architectural changes during contraction, which typically requires multi-stack acquisitions thereby increasing the total acquisition time in direct proportion to the number of stacks acquired. Hence, to be able to obtain information about architectural properties in a whole muscle, during a contraction, a different approach is required.

Planar ultrasonography has been used in dynamic settings to characterize changes in architectural properties with a high temporal resolution, but lacks the coverage required to visualize the whole muscle (34–37) and limited soft tissue contrast. Recent work showed that high resolution displacement fields derived from MR image registration were able to accurately quantify tissue displacement and strain in skeletal muscle during isometric contractions (38, 39). Based on these findings, we hypothesize that we can use displacement fields derived from registration of high-resolution anatomical images in different ankle positions to transform muscle fiber-tracts from a plantarflexed to a dorsiflexed ankle position, without significant differences in the observed mean architectural properties from those directly observed in the dorsiflexed position. If so, this would provide critical proof of the concept that a registration-based approach could be similarly used to transform the DTI-observed skeletal muscle architecture patterns due to contraction. The accuracy of the displacement fields, and thus the transformation of muscle fiber-tracts, depends highly on the quality of the registration. Therefore, we first optimized the registration parameters of the anatomical images; thereafter used interpolation to transform fiber-tracts from a plantarflexed to a dorsi-flexed ankle position; and finally determined whether the original tracts’ mean architectural properties differed from those of the transformed tracts.

## Materials and methods

### Study participants

Seven healthy volunteers (5 men; age: 25.1 ± 2.7 yrs. with range 23 – 31 yrs.) provided written informed consent to participate in this IRB-approved study. The written consent was obtained between 01-09-2019 and 01-11-2019. The inclusion criteria were healthy men and woman in the age range of 18 - 40 years. Exclusion criteria included the inability to provide informed consent, MRI contraindications, claustrophobia, physician-diagnosed muscle disease or other disease affecting muscle function, an inability to exercise safely, drug and alcohol use to the point of intoxication more than twice a week, and smoking tobacco. All participants were asked to abstain from alcohol (24 hours prior to the examination), caffeine (6 hours prior to the examination), and moderate or intense exercise (24 hours prior to the examination).

### MR examination

MR datasets were acquired in the right lower leg on a 3 Tesla MR System (Philips Elition; Best, the Netherlands) using a 16-element receiver coil (anterior) and the 10-element receiver coil built into the patient table (posterior). The participants were positioned supine, feet-first in the MR scanner with the right leg as close as possible to the centre of the bore. The foot was placed in an MR compatible exercise device which could be fixed in ankle angles ranging from +25° degrees plantarflexion to −15° degrees dorsiflexion. The anterior coil was placed on top of the legs and supported with foam pillows and fixation bands to ensure that the coil covered the full lower leg, did not compress the muscle, and did not move during the change of foot position from plantarflexion (+20°) to dorsiflexion (−10°). MRI data were first acquired with the foot passively held at +20° and again after the foot was passively rotated to −10°. The MR examination consisted of:

I. Chemical shift-based water-fat separation scan for anatomical reference and muscle and aponeurosis segmentation. (3D FFE; mDixon-Quant; TR/TE/ΔTE/FA, 210ms/1.01ms/0.96ms/3°; voxel size, 1×1×1.75mm^3^; number of excitations (N_EX_), 1; acquired matrix, 192×192; number of slices, 176; no slice gap; SENSE, 2; and scan duration, 36.2 seconds)
II. Two diffusion-weighted acquisition to assess muscle fascicle architecture (SE-EPI; TR/TE, 4800ms/53ms; 24 directions; b-values, 0 and 450 s/mm^2^; recon voxel size, 1×1×7mm^3^; N_EX_, 6; acquired/reconstructed matrices, 96×96/192×192; number of slices, 24; no slice gap, SENSE, 1.7; combination of three fat suppression techniques: SPectral Adiabatic Inversion Recovery (SPAIR) and Slice Selected Gradient Reversal (SSGR) for the aliphatic lipid peaks and a SPIR pulse for the olefinic peak (40); scan duration, 725 seconds). The DTI data were acquired in two transverse stacks with 28 mm overlap, fully covering 308 mm in the superior-inferior direction with a field of view (FOV) of 192×192 mm^2^.

For each type of sequence, the superior end of the slice stack was positioned at the level of the tibia plateau and the image stacks were rotated to align their midpoint with the tibia bone.

### Data-analysis

All data-analyses were performed in MATLAB R2019 (Mathworks) using the publicly available MuscleDTI_Toolbox (41) and additional custom written scripts. The full workflow of the study is shown in Figure 1.

**Figure 1.**
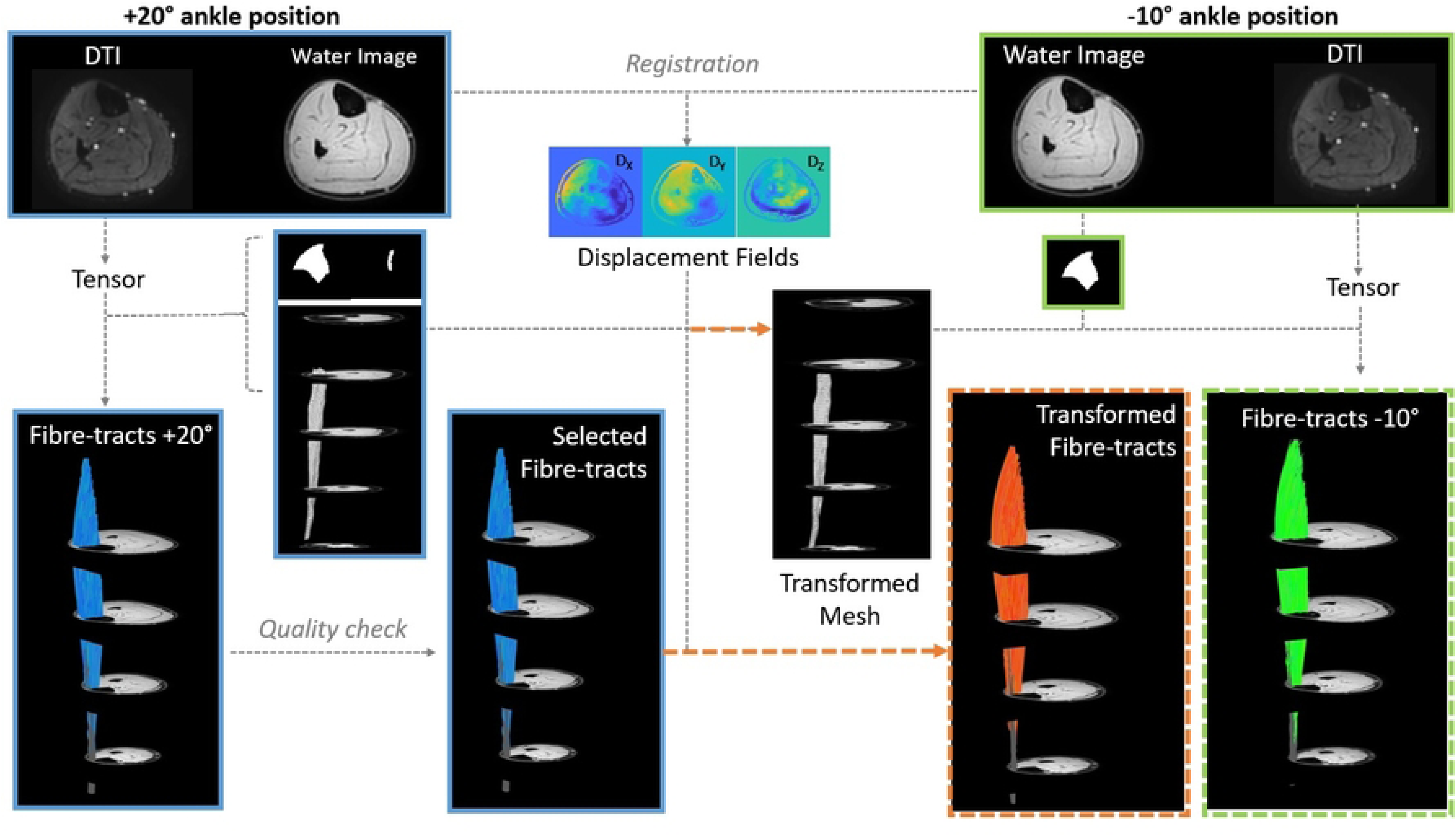
**Overview of the analysis workflow.** MR data were acquired in two ankle positions, +20° (blue outline) and −10° (green outline). Anatomical images were manually segmented to derive masks for the TA aponeurosis and TA muscle (binarized images). After the tensor calculation, fiber tracking was performed. A quality check was used to exclude obviously erroneous results (resulting in the selected fibers). Displacement fields were used to transform fiber-tracts and the aponeurosis mesh from +20° to −10° ankle position (orange arrows). The transformed aponeurosis is used to initiate tracking in the −10° position to allow direct comparison between transformed (orange dotted outline) and original fiber-tracts (green dotted outline).

### VOIs and aponeurosis mesh

For each foot position, masks of the Tibialis Anterior (TA) boundaries and aponeurosis were manually drawn in ITK-snap (version 3.8.0; www.itksnap.org) (42) on the high resolution mDixon-Quant water image. The manually segmented aponeurosis masks were converted to a low-resolution seed-point mesh (mesh size: 30×20) using *define_roi*( ). An additional high-resolution seed-point mesh (size 100×50) was also defined for visualizing the outcomes of the optimized method.

### Optimization of anatomical image registration

The high-resolution anatomical scans from the +20° ankle position were registered to the anatomical scan from the −10° position using MATLAB’s *imregdemons*( ) function (2000 iterations, 4 pyramid levels and AccumulatedFieldSmoothing of 1.0; Fig. 1). This function uses a 3D registration based on a Demons algorithm (43, 44), resulting in displacement fields for the X, Y and Z directions. A variety of registration inputs were explored, including various contrasts (*2 settings; water image and out-of-phase images*); slice thickness (*3 settings; 7 mm, 3.5 mm and 1.75 mm*); normalization (*2 settings; normalized images and non-normalized images*); and masking options (*2 settings; full mask lower leg; mask for the TA muscle)*. Image normalization was performed by dividing the signal intensities in the original water or out-of-phase image by the maximal signal intensity in the image slice. Masking of the full lower leg was based on an empirically determined signal intensity threshold. For the TA-only mask, the manually segmented boundary mask was used.

The registration displacement fields were smoothed using a Savitzky-Golay filter with a polynomial order of 2 and frame length of 2, as implemented in the MATLAB function *sgolay*( ). This registration field was used to transform the muscle boundary mask and the seed-point mesh from +20° foot position to the −10⁰ position by 3D interpolation (MATLAB’s function *interp3*) to the −10° foot position; see Fig. 1. For each set of registration inputs, the outcomes were assessed by calculating the Sorensen Dice Similarity Coefficient (DSC) for the muscle mask (DSC_mask_) and aponeurosis (DSC_apo_) mask and the average Hausdorff distance (D_H_) and Euclidean distance (D_E_) between corresponding points on the aponeurosis mesh. The optimal registration approach was defined as the method with the best ordinal ranking across all outcome assessments.

### Pre-processing of diffusion images

The diffusion images were registered to the scanner-reconstructed water images of the Dixon series using the *imregdemons*( ) function. Then the diffusion data were denoised using an anisotropic smoothing algorithm (45, 46) using a noise level of 5% (47) The diffusion tensor was estimated in each voxel from the denoised diffusion data using weighted least squares fitting(41).

### Fiber-tracking

The diffusion tensor field, seed-point mesh, and muscle boundary mask were used as inputs for fiber-tracking. Fiber-tracts were initiated from points along the seed-point mesh and propagated using 4^th^-order Runge-Kutta integration of the principal eigenvector at a step size of 1.0 pixel-width. Stopping criteria included either 1) reaching the muscle mask boundary or 2) if two consecutive data points had a Fractional Anisotropy value <0.1 or >0.4 or a trajectory angle between consecutive tracking steps of >30°. Three sets of fiber-tracts were generated:

1. Fiber-tracts were generated for the +20° foot position after initiation from the seed-point mesh originally defined for this position (Original +20°).
2. The seed-point mesh was transformed from the +20° position to the −10° position. Fiber-tracts for the −10° position were then generated from the transformed mesh (Original −10°). The transformed mesh was used to enable direct comparison between the transformed and original datasets.
3. The fiber-tracts from the +20° position were transformed to their predicted locations at a foot angle of −10° (Transformed −10°). Transformation was performed as described above.

### Architectural quantification and goodness filtering

Prior to architectural quantification of the fiber-tracts, the row, column and slice positions were smoothed using 3^rd^ order polynomial fitting (48, 49). Architectural tract properties, including pennation angle (θ; (7)), fiber-tract length (L_FT_), and curvature (*κ*; (48)), were quantified for each tract in all subjects using the *fiber_quantifier*( ) function. Pennation angle was defined as the complement to the angle formed by the normal vector to the aponeurosis at the seed point and the position vectors. The tract length was calculated by summing the inter-point distances, and the curvature was calculated using a discrete implementation of the Frenet-Serret equations.The *fiber_goodness*( ) function was used to exclude fiber-tracts if their Z-positions did not increase monotonically; if their mean curvature was >40 m^-1^, their mean pennation angle was >40°, and/or their length was < 10 mm or > 98 mm; or if there were local outliers in any of these architectural properties. The maximum length was based on the average fascicle length plus two times the standard deviation reported in an ultrasonography study (54) In addition, the muscle volume (V_M_), mean (L_FT_), and θ were used to calculate the Physiological Cross-sectional Area (PCSA) as (50): PCSA = V_M_ * cos (θ) / L_FT_, using L_FT_ as a proxy for fascicle length.

### Fiber Similarity Measures

To compare the original and transformed −10⁰ fiber-tract outcomes, we determined the differences in the mean pennation angle (Δθ), fiber-tract length (ΔL_FT_) and curvature (Δ*<u>κ</u>*) for the deep compartment, superficial compartment, and whole muscle. Additionally, the similarity (S_i_) was calculated between the transformed and its corresponding original fiber (similar mesh starting point) using: S_i_ = R_CS_ · e ^(– D^_E_^/C)^ (8, 51), where R_CS_ is the corresponding segment ratio, D_E_ is the mean Euclidean distance between corresponding points, and C is a weighting factor (here set to 1.50 mm). We used the median S_i_ to characterize this property on a whole-muscle level. As exploratory analyses, we also compared the differences in architectural properties and S_i_ at the individual fiber-tract level.

### Statistical Analysis

All statistical analysis were performed using SPSS (IBM SPSS Statistics 28.01.1.1, Armonk, NY: IBM Corp). Data were checked for normality using a Shapiro – Wilk test. Paired *t*-tests or the non-parametric version of the *t-*test were used to evaluate the difference in mean architectural properties (θ, L_FT_, *κ,* and PCSA) between the original and transformed fiber-tracts in the deep compartment, superficial compartment, and whole muscle. Pearson correlation and Bland-Altman analyses were used to exploratory evaluate architectural properties on the fiber-tract level. Furthermore, Spearman correlations were used to explore the relation between the registration outcome measures (DSC_mask_, DSC_apo_, D_H_ and D_E_) and the similarity measures (Δθ, ΔL_FT_, Δ*<u>κ</u>* and S_i_).

## Results

### Data quality

All MR datasets were complete and without observable fat and/or motion artefacts.

### Registration approaches

The DSC, D_H_ and D_E_ for the muscle mask, aponeurosis mask and aponeurosis mesh using the different registration approaches are shown in Table 1. Overall, the non-normalized out-of-phase (TE= 1.01ms) images with a slice thickness of 3.5 mm resulted in the lowest ordinal scale and were selected to transform the aponeurosis mesh and fiber-tracts. One of the datasets did not register correctly, as assessed by visual inspection by MH and low DSC values for the muscle and the aponeurosis masks. Thus, for the comparison of architectural properties and correlation analysis we excluded this dataset.

**Table 1.**
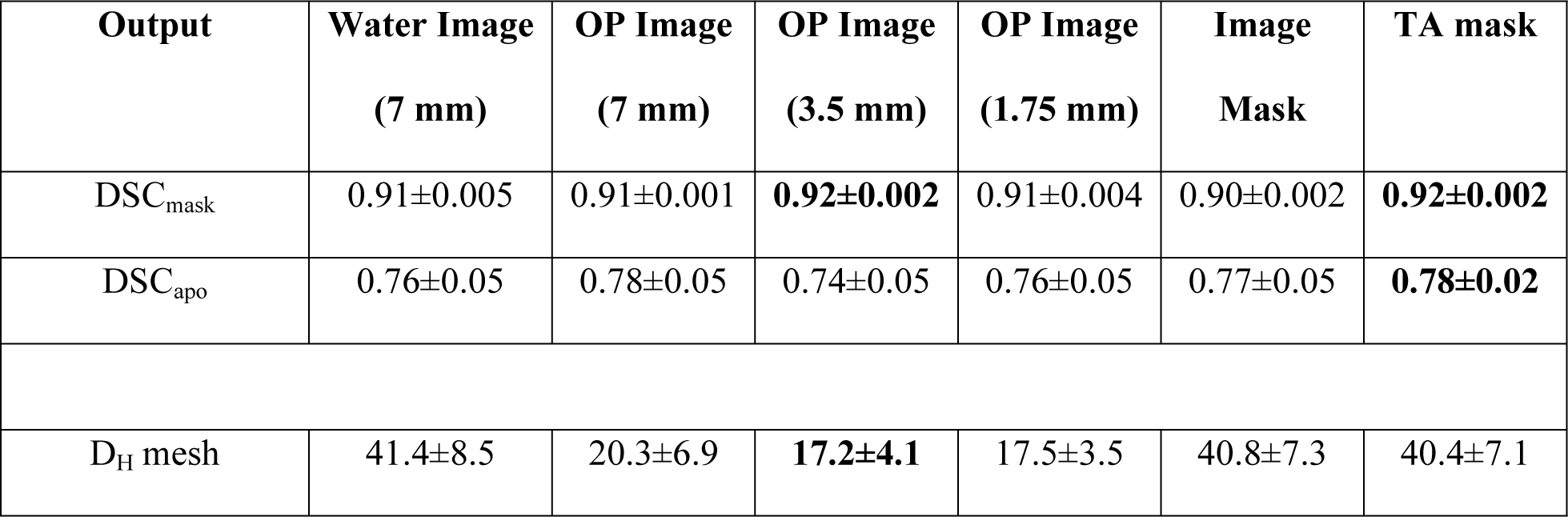
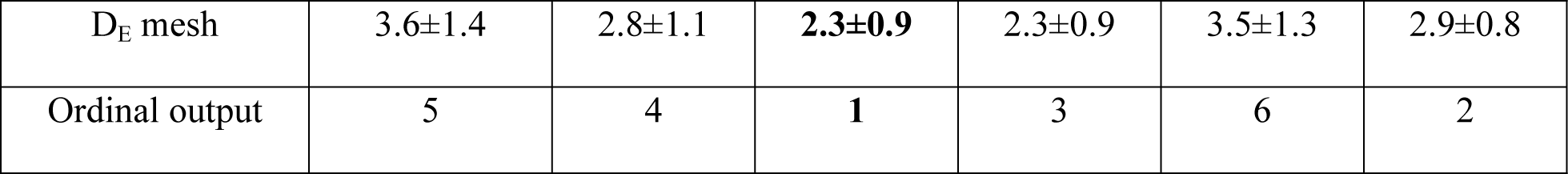
Evaluation of the registration approaches. **Table legend 1**: Data table showing the average and standard deviation of the DSC, Hausdorff distance (D_H_), Euclidean distance (D_E_) and overall ordinal grading for the muscle mask, aponeurosis mask and aponeurosis mesh using a selection of the different registration strategies. In bold text is shown the best approach per category. Note the overall best strategy: Non-normalized Out-of-phase images with a slice thickness of 3.5mm.

**Table 2.** Architectural properties of the original fascicles in the −10° and +20° ankle position and the transformed fascicles. **Table legend 2**: Data table showing the average and standard deviation pennation angle, fiber-tract length and curvature values for the original fiber-tracts in the dorsiflexed −10° ankle position, plantarflexed +20° ankle position and the transformed fiber-tracts.

| Outcome measure | Ankle position | Full TA muscle | Deep compartment | Superficial compartment |
| --- | --- | --- | --- | --- |
| Angle (°) | Original +20° | 6.3±1.0° | 6.4 ±1.5° | 6.5 ±1.2° |
|  | Original -10° | 9.1±1.4° | 10.7 ±3.3 | 10.0 ±3.5° |
|  | Transformed -10° | 7.3 ±1.7° | 5.9 ±1.2 | 7.8 ±1.4° |
| **Length (mm)** | **Original +20°** | 51.5 ±4.51 | 56.4 ±13 | 59.2± 8.8 |
|  | **Original -10°** | 49.9 ±4.25 | 52.3 ±8.6 | 47.7 ±7.6 |
|  | **Transformed -10°** | 48.3 ±10.8 | 50.7 ±13.7 | 43.9 ±9.8 |
| **Curvature (m<sup>-1</sup>)** | **Original +20°** | 7.2 ±0.7 | 7.2 ±1.6 | 6.9 ±0.6 |
|  | **Original -10°** | 8.7 ±1.4 | 7.5 ±3.0 | 9.8 ±2.1 |
|  | **Transformed -10°** | 9.8 ±3.2 | 9.8 ±4.5 | 9.8 ±2.1 |

### Architectural properties

Representative original and transformed datasets, using the high-resolution seed-point mesh definition and the optimized registration criteria from Table 1, are shown in Figure 2. Data were normally distributed and analysed using a paired t-test. No significant differences were detected between the original and transformed fiber-tracts for the mean values of θ, L_FT_, and *κ* in the deep compartment (*p*-value θ: 0.64, L_FT_: 0.74, and *κ*: 0.39), superficial compartment (*p*-value θ: 0.10, L_FT_: 0.44, and *κ*: 0.92), and whole TA muscle (*p-*value θ: 0.13, L_FT_: 0.72, and *κ*: 0.52)) (Figure 3). No significant differences were detected in PCSA between the original and transformed fiber-tracts in the whole TA muscle (*p-*value: 0.454). On an individual fiber-tract level variations were observed in the agreement in pennation angle and fiber-tract length between the original and transformed fiber’s for both a high and a low similarity dataset (Supplemental Figure 1). A significantly higher pennation angle (*p* = 0.004) was found in −10° ankle position compared to +20°, while no differences were detected in fiber-tract length and curvature between the two ankle positions (*p* = 0.54) (Figure 4).

**Figure 2.**
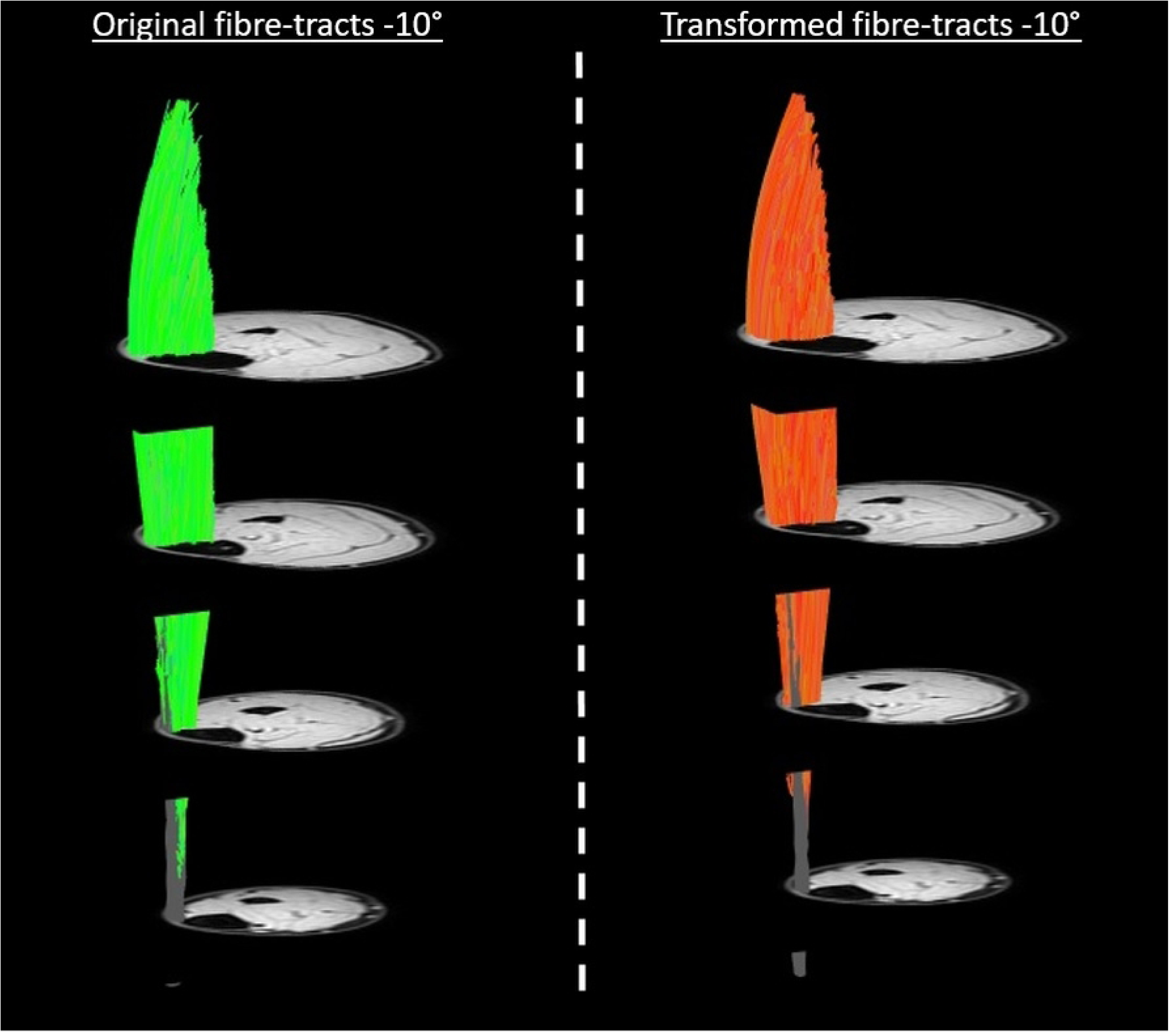
**Original and transformed fiber-tracts**. The original fiber-tracts (green) and aponeurosis mesh are shown on the left side and the corresponding transformed fiber-tracts (orange) and aponeurosis mesh on the right side for a representative dataset. For visualization purposes we used a high-density mesh (size 100×50).

**Figure 3.**
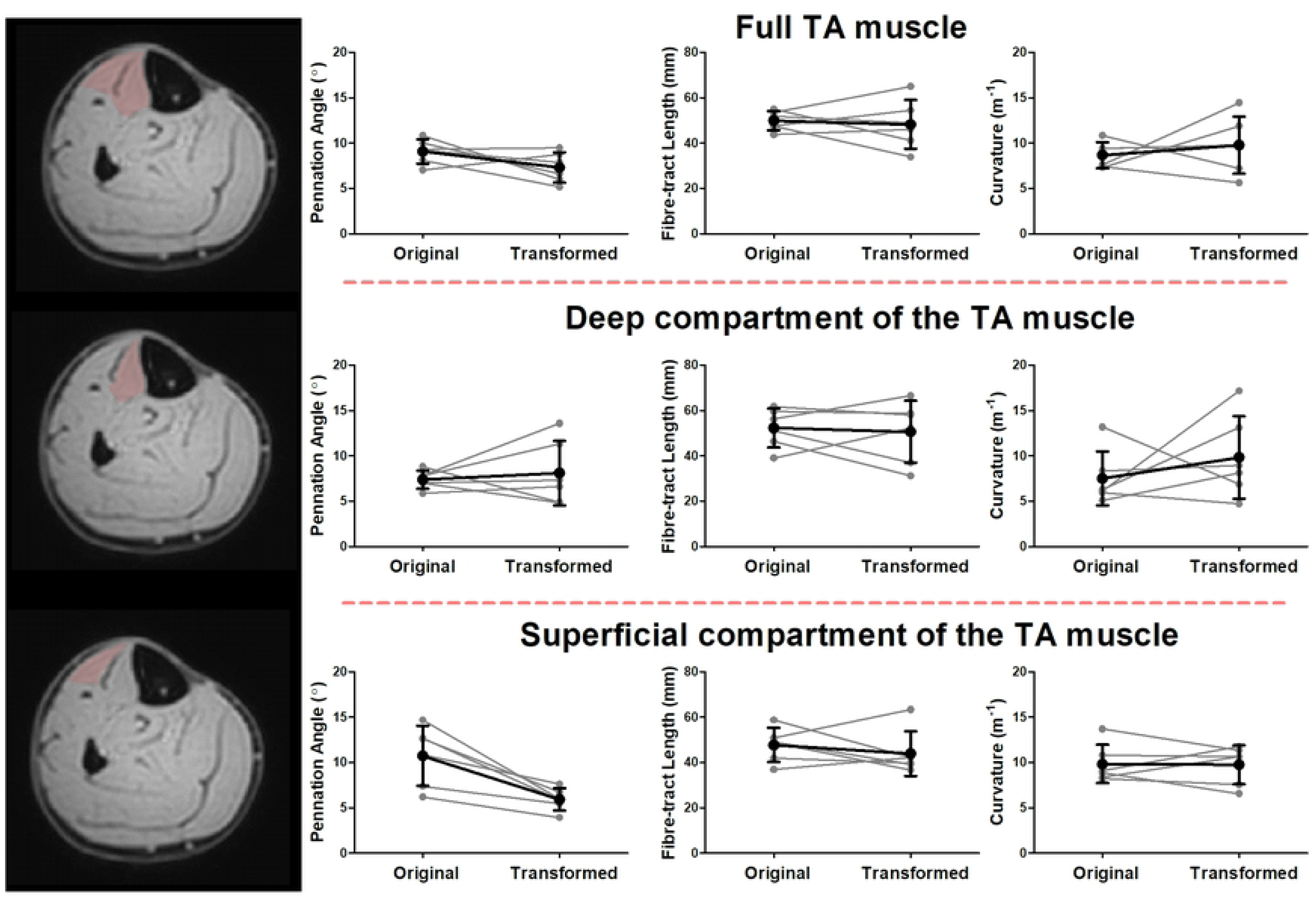
**Architectural properties of the original and transformed fibers.** Line plots showing the pennation angle (left), fascicle length (middle) and fascicle curvature for the original and transformed fiber-tracts in the full TA muscle (top row), the deep compartment of the TA muscle (middle row) and the superficial compartment of the TA muscle (bottom row). In black the mean and standard deviation for all the subjects and in light grey the individual subjects. Significant differences are indicated with an asterisk.

**Figure 4.**
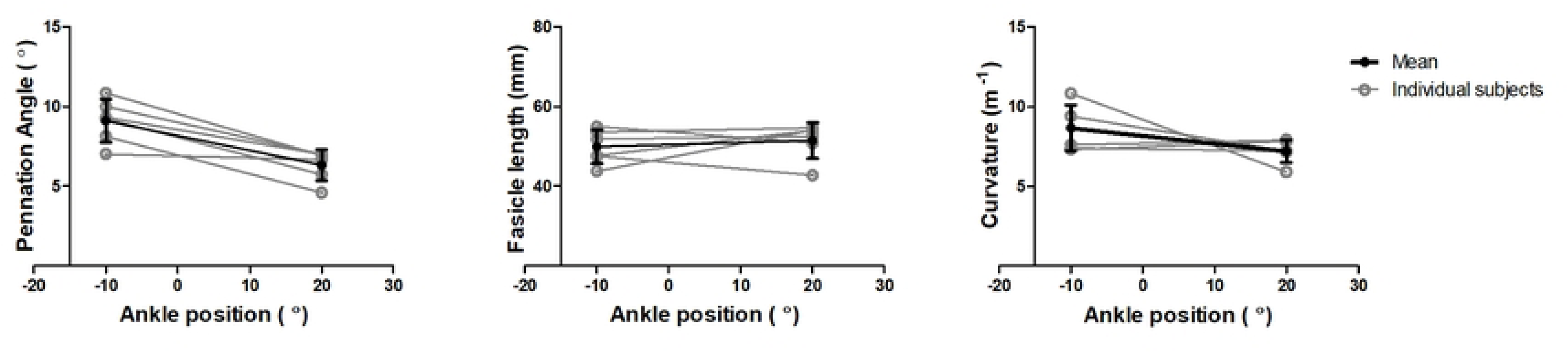
**Pennation angle and fascicle lengths of the original fibers in the two foot positions** Line plots showing the pennation angle (left) and fascicle length (middle) and the curvature (right) for the original fiber-tracts in −10° and +20° ankle position in the full TA muscle. In black symbols are shown the mean and standard deviation for all the subjects and in light grey symbols are shown the individual subjects’ data. Significant differences are indicated with an asterisk.

### Registration quality measures in relation to fiber similarity measures

On a whole muscle basis, no significant correlations were observed between the similarity (S_i_) or registration quality measures (D_H_ (r=-0.41; p=0.42), D_E_ (r=0.08; p=0.91), DSC_apo_ (r=0.64; p=0.18) and DSC_mask_ (r=-0.65; p=0.18) and Δθ (r=-0.42; p=0.42), ΔL_FT_ (r=-0.26; p=0.66) (Figure 5.) or Δ*κ*. Similarly, no clear pattern was evident between Δθ and ΔL_FT_ and the registration quality measures (Spearman R absolute range = 0.03-0.67; *p* > 0.18) (Supplemental Figures 2 and 3). Within subjects, the similarity values ranged widely (between 0.03-0.77). Typical example fiber-tracts for a low and high similarity cases are shown in Figure 6. The average per-subject similarity value ranged between 0.14 and 0.31 across the sample. We did not find a clear pattern for the comparison of S_i_ and Δθ, ΔL_FT_ on a fiber-tract basis for both the highest and lowest averaged similarity score (Supplemental Figure 4).

**Figure 5.**
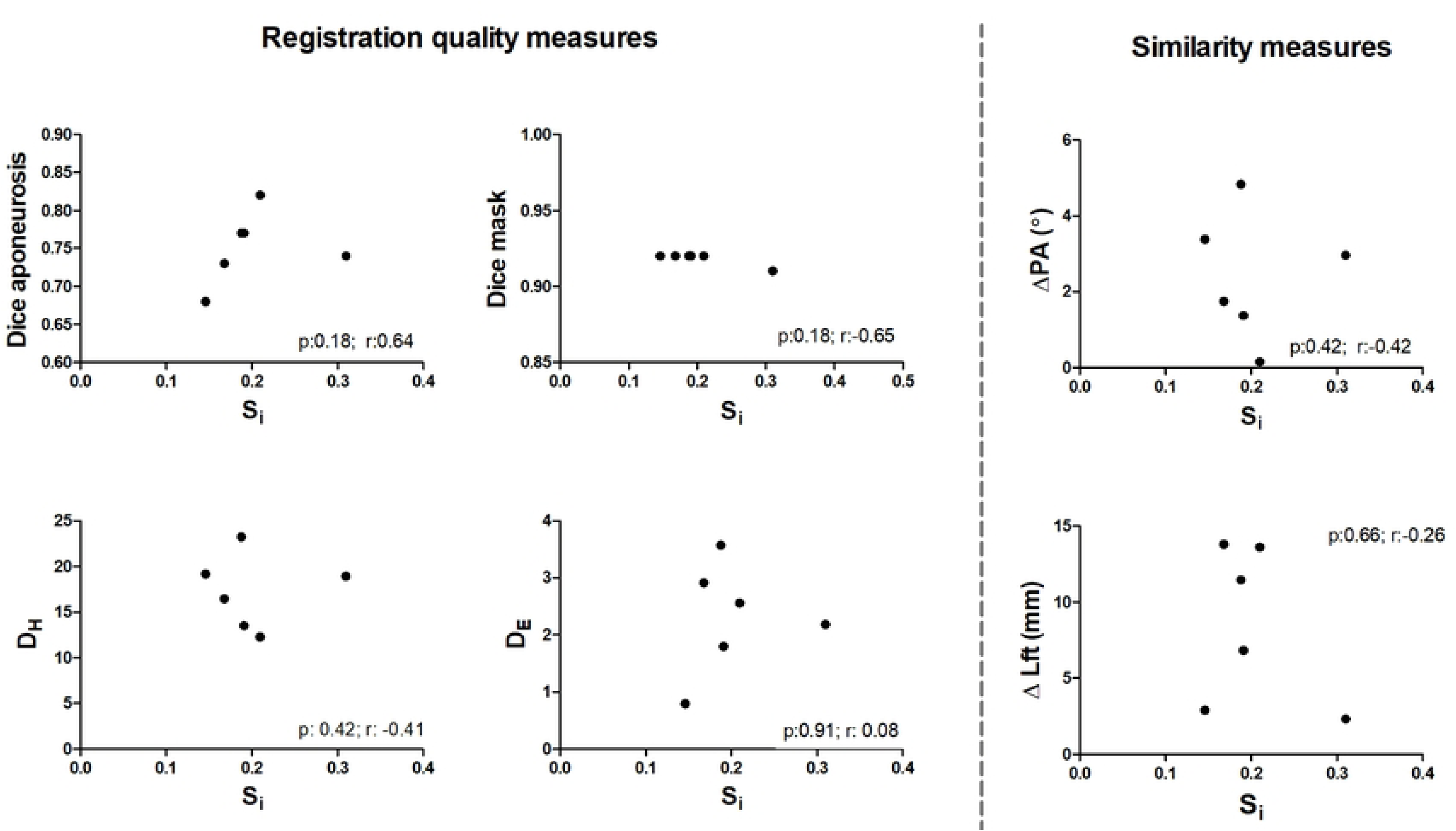
**Correlation analysis between the Similarity index, similarity measures and registration measures.** Dot plots showing the correlation between the tract similarity measure and registration quality measures. The black dots represent the individual participants. D_H_: Hausdorf Distance, D_E_: Euclidean Distance; Δθ: the difference in pennation angle between the original and transformed fiber-tracts, ΔL_FT_: the difference in length between the original and transformed fiber-tracts.

**Figure 6.**
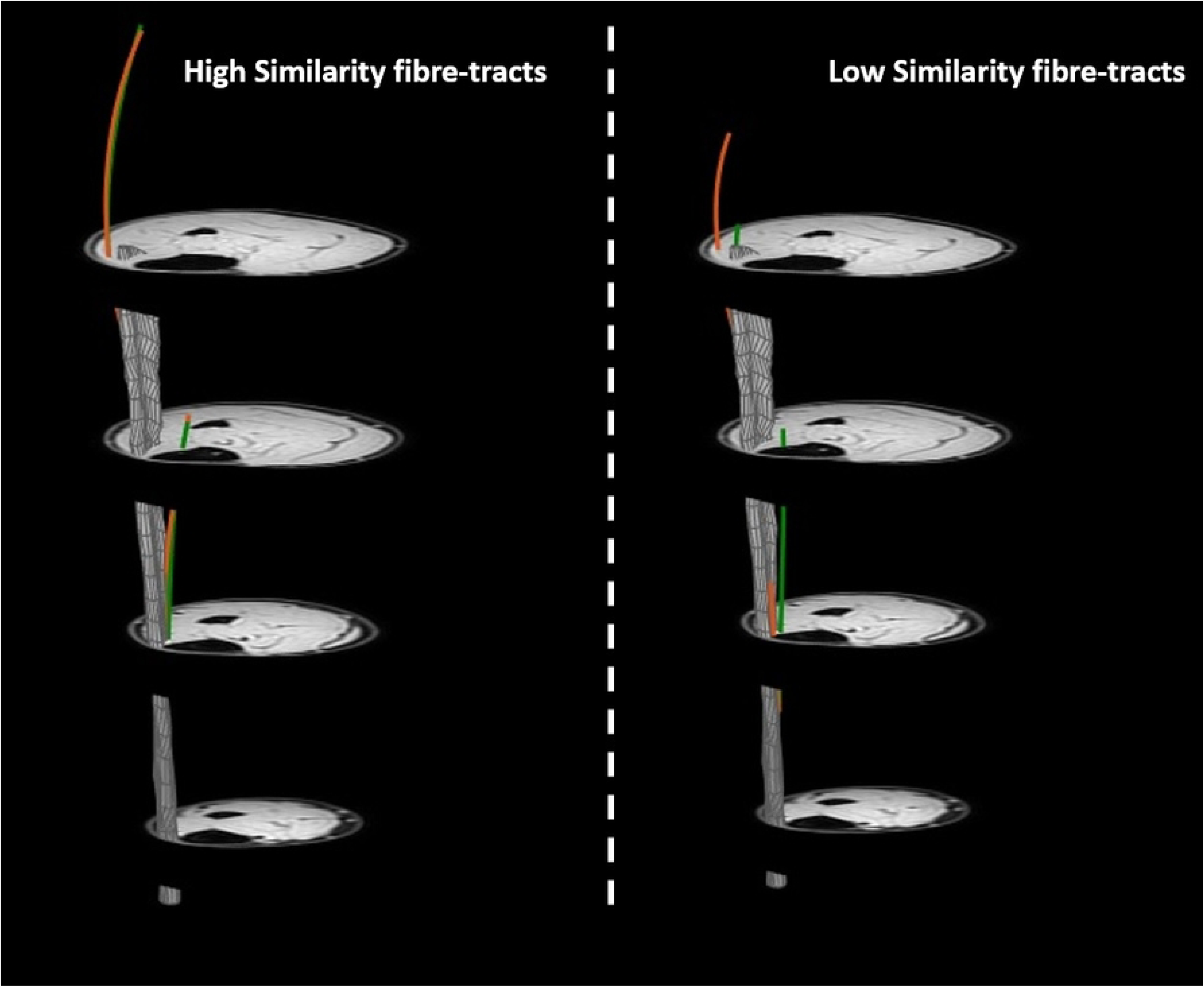
Visualization of the transformation of high and low similarity fiber-tracts. Visualization of variations in the quality of transforming muscle architecture by highlighting two individual fiber-tracts (one with a more proximal aponeurosis attachment and one with a more distal aponeurosis attachment). In green the original fiber-tracts and in orange the corresponding transformed fiber-tracts for one representative subject. The image on the left shows original and transformed fiber-tracts with high similarity (S_i_ =0.53 & 0.55) and the image on the right displays original and transformed fiber-tracts with low similarity(S_i_ =0.03 & 0.07).

## Discussion

We showed that DTI-determined muscle fiber-tracts can be transformed from a plantarflexed ankle position to the dorsiflexed ankle position using registration of high-resolution anatomical images. On the whole-muscle and whole-compartment scales, the original and transformed fiber-tracts did not differ in architectural characteristics, i.e. mean length, pennation angle, curvature, and PCSA. This finding emphasizes the potential of this approach for analysing compartmental or whole-muscle variations in architectural properties during passive movements about a joint. However, there was variability in the quality of the outcomes across participants and across fiber-tracts within a participant. The applicability of this method therefore depends on the intended application of the data.

To our knowledge, this is the first study exploring the use of high-resolution displacement fields to transform DTI-determined muscle fiber-tracts between joint angles. The original and the corresponding transformed fiber-tracts showed similar mean architectural properties at the whole-muscle and compartmental scales. These properties were also in the same range as pennation angles and fascicle lengths previously reported in the TA muscle using DTI (7, 15, 16, 52–54), ultrasonography (55–58), and cadaver (59, 60) studies. When comparing the original and transformed tracts from the −10⁰ position, we did observe compartment-specific variations in the fiber-tract lengths, but these did not rise to the level of statistical significance.

Besides non-invasively reflecting anatomy, muscle fiber-tract architecture is also used to predict function (3, 61) and to detect changes in architectural properties between conditions. Indeed, the PCSA (62) is considered one of the key predictors of muscle force production; therefore, not finding changes in PCSA between the transformed and original fiber-tracts is very relevant with respect to the use of this method in musculoskeletal modelling (25–30). Additionally, we observed a smaller pennation angle in the full TA in the plantarflexed (+20°) compared to dorsiflexed (−10°) ankle position, which is consistent with physiological predictions based on the relative compliance of resting muscle vs. tendinous structures and previous observations (55, 63, 64). Interestingly, we did not observe a clear pattern for fiber-tract length on a group-level basis, and both longer and shorter fiber-tracts were seen across participants. This was contrary to both physiologically based predictions and prior work on the full TA muscle that showed a reduced fascicle length in dorsiflexed position (−15°) compared to plantarflexed (+30°) ankle position (24, 65). Possible causes for the shorter fiber-tracts are muscle tendon complex lengthening, or some inconsistency or error in fiber-tracking due to a small mismatch in registration between the DTI and anatomical data.

The quality of the transformation of fiber-tracts highly depends on the performance of the registration, in this case using the Demons algorithm. Previously, the Demons algorithm has been used and validated to accurately quantify displacement and deformations in skeletal muscle tissue (38, 66), pelvic floor, lung and cortical bone tissue (67, 68). In addition, the reliability of this algorithm has been tested by imposing known deformations to an image set (38) with the measured strains being conservative estimators of the local deformations. To establish the quality of fiber-tract transformations, we evaluated a variety of registration inputs to optimize displacement fields derived from the Demons algorithm and showed that the out–of–phase (OP) images with an intermediate slice thickness led to the best registration result. This may be due to the higher internal contrast between anatomic structures in these images compared to the water-only images, the lower leg mask, and muscle mask. The difference in registration quality between the three slice thicknesses was minimal, which may indicate that these slice thicknesses are sufficient to sample any foot-head variations in structure, at least in the case of this muscle in healthy adults. The in-plane resolution of the anatomical images used for registration was similar to the resolution of the DTI data. Future work could explore if higher in-plane resolution could benefit the registration further.

In addition to comparing the compartmental and whole-muscle architectural properties of the entire dataset, similarity and architectural measures were evaluated at the fiber-tract level. The Bland-Altman analysis clearly visualized variations in similarity in architectural properties on an individual fiber-tract level, independent of a high or low similarity analysis. This finding is consistent with the correlation analysis which displayed clearly that tracts with both high and low similarities could have very similar architectural properties and that tract pairs with low similarity values could exhibit either high or low levels agreement among their architectural properties (Supplemental Figure 4). This suggests that the similarity index used here does not adequately assess fiber-tract likeness. This may be because the similarity measure accounts for both the amount of length overlap of two tracts and the distance between their corresponding points, allowing both adjacent fiber-tracts and fiber-tracts that lay further apart to have the same similarity scores (8, 51, 69). Also, our registration quality measures did not show convincing correlations with our fiber similarity measures. This was contrary to our expectations, since we optimized according to these measures; however, this evaluation may have been impacted by the low number of datasets, the imperfect nature of this similarity index as an outcome measure, and the small dynamic range of some of the registration outcome parameters. The problem of low dynamic range is exacerbated by excluding the dataset that did not register properly. In the next development phase of the registration-based approach to transforming fiber-tracts, it may be useful to expand the calculation of the similarity measure to also include differences in architecture between the tracts or include other structural features as terms in the similarity calculation.

The end goal of this approach is to explore muscle architecture during dynamic contractions. To address that goal, DTI data would be acquired in relaxed position and then transformed to its contracted state using anatomical images acquired at rest and during contraction. This approach necessarily rules out the possibility to compare fiber-tracts according to the similarity measure or the observed architectural properties, making validation less straightforward. From pilot data, we know that smaller displacements occur from relaxed to contracted state in a fixed ankle angle isometric contraction, suggesting that registration of these type of datasets should be less challenging. Furthermore, the displacement fields derived from the registration have also been used to quantify strain values in muscle tissue; which showed to correspond well with values measured with other techniques (MR-tagging and PC-MRI). The amount of correspondence between strain values derived from displacement fields and measured with other MR techniques during active contractions could possibly function as justification in active situations. All combined, this study emphasizes the potential of a registration-based approach for observing whole-muscle or compartment changes in architectural properties during isometric muscle contractions.

Our study has several limitations. First, it is a proof-of-concept study, in a limited number of young, healthy participants, which makes detecting meaningful differences challenging. Consequently, no difference between the original and transformed fiber-tracts does not automatically indicate the accuracy of this registration-based approach; rather, this work primarily showcases its potential. Further, the translatability to other populations (such as aging and neuromuscular diseases) should be explored further. However, some of the pathological features known to characterize aging and diseased muscle, i.e. the replacement of muscle tissue by fat, result in more contrast in the images, potentially benefitting the registration and the transformation of muscle fiber-tracts. Additionally, how well the fiber-tracts can be transformed depends highly on the performance of the registration. Here we only evaluated a narrow band of data for which the registration worked well and therefore may have overestimated the performance of the approach. Despite the relatively high DSC indices and low D_H_ and D_E_ we found using our registration approach, alternative or novel registration strategies or approaches could improve the outcomes from this study even further. We also note that the method did not perform as well at the fiber-tract level as it did at the compartmental and whole-muscle levels, which again motivates exploration of alternative or novel registration strategies. Lastly, we used the fiber-tracts generated in −10° ankle position with the transformed mesh as ground truth. However, noise and other image quality issues will affect the performance of DTI tractography (65, 70, 71), perhaps leading to an underestimation of the performance of our approach.

In summary, we showed that muscle fiber-tract architecture from one ankle position can be transformed in the other ankle position using registration derived displacement fields. Whole-muscle architectural characteristics, i.e. fiber-tract length, pennation angle, curvature, and physiological cross-sectional area of the original and transformed fiber-tracts did not differ significantly on a group-level basis. Consequently, this approach to transform muscle architecture is very promising, and our next step will be to evaluate if muscle architecture can be reconstructed in a contracted state using a similar approach.

## Acknowledgements

The authors thank Hannah Kilpatrick and Mark George for their technical assistance. BD acknowledges NIH grants NIH R01 AR073831 and NIH S10 OD021771. The sponsor did not play a role in the study design, data collection and analysis, decision to publish or preparation of the manuscript.

## Competing Interest Statement

The authors have declared that no competing interests exist.

## Data Availability Statement

The fat/water MRI and unprocessed DTI data are available to any qualified investigator at an academic institution or private research organization who has current human subjects research ethics training certification and can describe a scientific use for the data. Interested parties can contact the corresponding author to establish a Data Use Agreement.

---

**Supplemental Figure 1.**
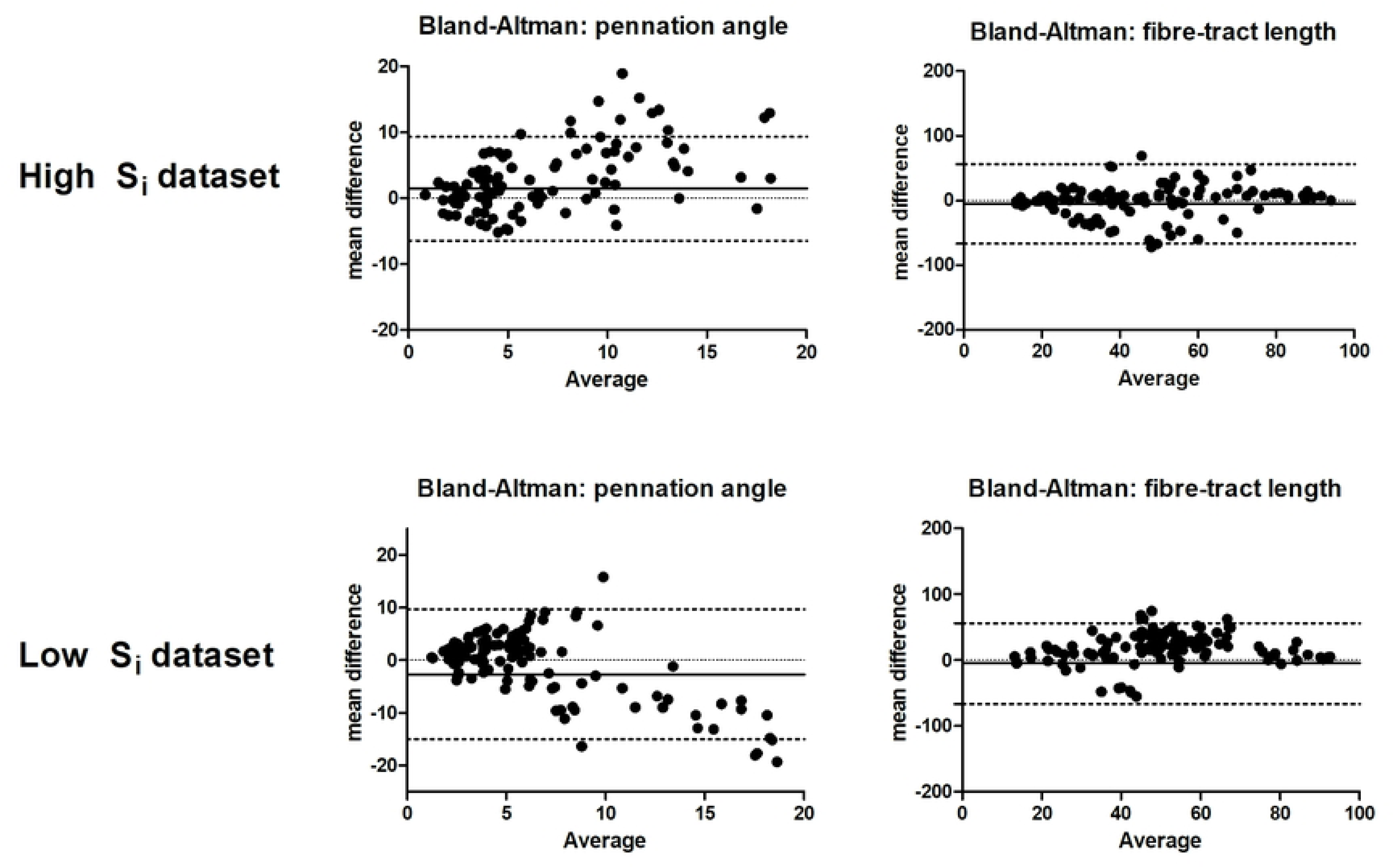
Bland-Altman Plots showing the similarity between the original and transformed fiber-tracts for pennation angle (Δθ) and fiber-tract length (ΔL_FT_), on a fiber tract level, for high similarity (left) and low similarity (right) dataset. Note differences in Y-axis scales between the left and right panels.

**Supplemental figure 2.**
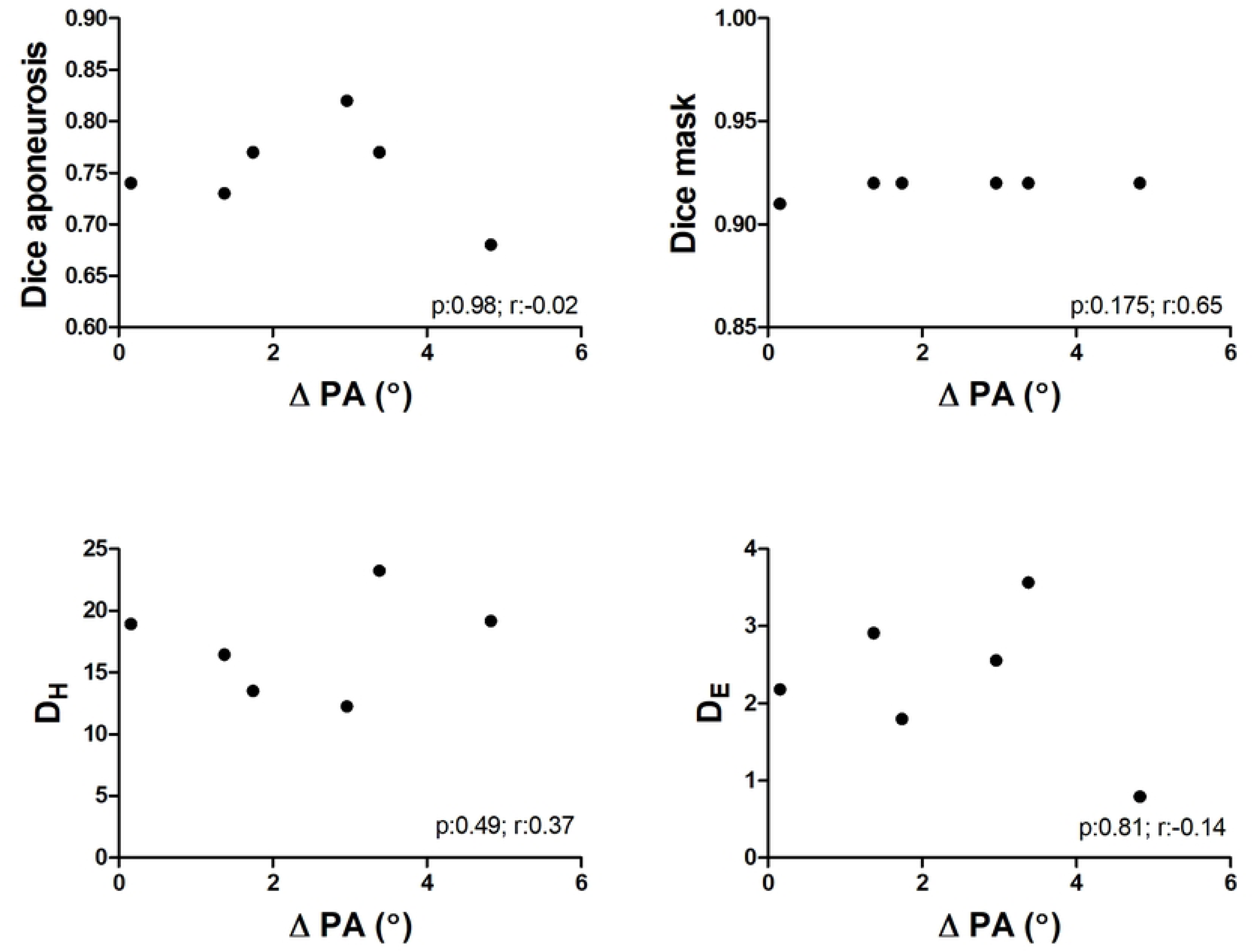
Dot plots displaying the correlation between the registration quality measures and the difference in pennation angle (Δθ) between the original and transformed fiber-tracts for each of the participants (black dots).

**Supplemental figure 3.**
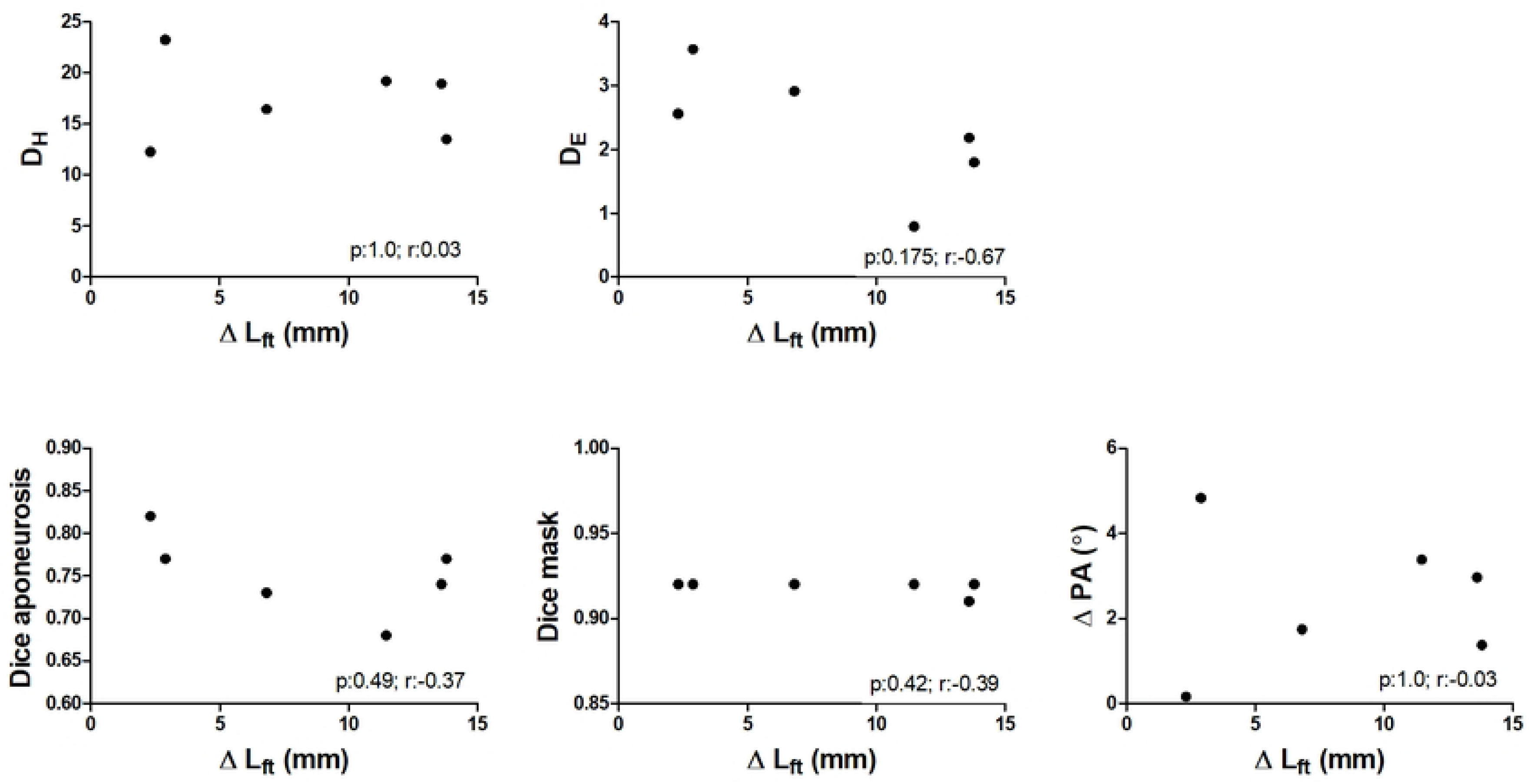
Dot plots displaying the correlation between registration quality measures and the difference in fiber-tract length (ΔL_FT_) between the original and transformed fiber-tracts for each of the participants (black dots).

**Supplemental Figure 4.**
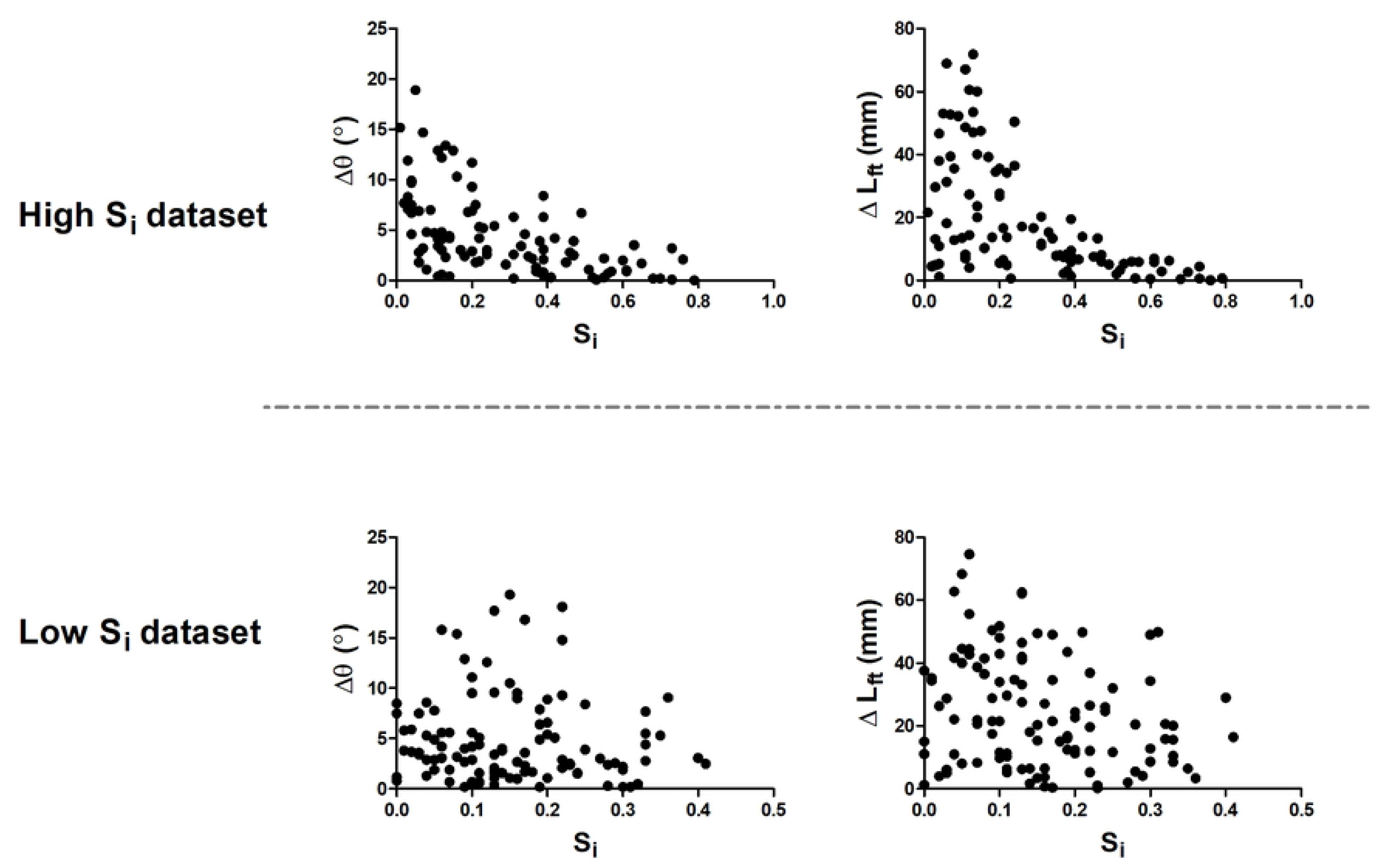
Scatterplots showing the relation between the similarity (S_i_) and the difference in pennation angle (Δθ) and fiber-tract length (ΔL_FT_) between the original and transformed fiber-tracts for each of the individual fiber-tracts for the dataset with the highest and lowest averaged similarity value.

## Notes

### Competing Interest Statement

The authors have declared no competing interest.

## References

1. Huijing PA. Architecture of the Human Gastrocnemius-Muscle and Some Functional Consequences. Acta Anat. 1985;123(2):101–7.

2. Sopher RS, Amis AA, Davies DC, Jeffers JR. The influence of muscle pennation angle and cross-sectional area on contact forces in the ankle joint. J Strain Anal Eng Des. 2017;52(1):12–23.

3. Lieber RL, Friden J. Functional and clinical significance of skeletal muscle architecture. Muscle Nerve. 2000;23(11):1647–66.

4. Basser PJ, Mattiello J, LeBihan D. MR diffusion tensor spectroscopy and imaging. Biophys J. 1994;66(1):259–67.

5. Basser PJ, Pierpaoli C. Microstructural and physiological features of tissues elucidated by quantitative-diffusion-tensor MRI. J Magn Reson B. 1996;111(3):209–19.

6. Noseworthy MD, Davis AD, Elzibak AH. Advanced MR imaging techniques for skeletal muscle evaluation. Semin Musculoskelet Radiol. 2010;14(2):257–68.

7. Lansdown DA, Ding Z, Wadington M, Hornberger JL, Damon BM. Quantitative diffusion tensor MRI-based fiber tracking of human skeletal muscle. J Appl Physiol (1985). 2007;103(2):673–81.

8. Damon BM, Ding Z, Anderson AW, Freyer AS, Gore JC. Validation of diffusion tensor MRI-based muscle fiber tracking. Magn Reson Med. 2002;48(1):97–104.

9. Heemskerk AM, Drost MR, van Bochove GS, van Oosterhout MF, Nicolay K, Strijkers GJ. DTI-based assessment of ischemia-reperfusion in mouse skeletal muscle. Magn Reson Med. 2006;56(2):272–81.

10. Galban CJ, Maderwald S, Uffmann K, de Greiff A, Ladd ME. Diffusive sensitivity to muscle architecture: a magnetic resonance diffusion tensor imaging study of the human calf. Eur J Appl Physiol. 2004;93(3):253–62.

11. Froeling M, Oudeman J, Strijkers GJ, Maas M, Drost MR, Nicolay K, Nederveen AJ. Muscle changes detected with diffusion-tensor imaging after long-distance running. Radiology. 2015;274(2):548–62.

12. Hooijmans MT, Damon BM, Froeling M, Versluis MJ, Burakiewicz J, Verschuuren JJ, et al. Evaluation of skeletal muscle DTI in patients with duchenne muscular dystrophy. NMR Biomed. 2015;28(11):1589–97.

13. Zaraiskaya T, Kumbhare D, Noseworthy MD. Diffusion tensor imaging in evaluation of human skeletal muscle injury. J Magn Reson Imaging. 2006;24(2):402–8.

14. Bolsterlee B, Finni T, D’Souza A, Eguchi J, Clarke EC, Herbert RD. Three-dimensional architecture of the whole human soleus muscle in vivo. PeerJ. 2018;6:e4610.

15. Foure A, Ogier AC, Le Troter A, Vilmen C, Feiweier T, Guye M, et al. Diffusion Properties and 3D Architecture of Human Lower Leg Muscles Assessed with Ultra-High-Field-Strength Diffusion-Tensor MR Imaging and Tractography: Reproducibility and Sensitivity to Sex Difference and Intramuscular Variability. Radiology. 2018;287(2):592–607.

16. Oudeman J, Mazzoli V, Marra MA, Nicolay K, Maas M, Verdonschot N, et al. A novel diffusion-tensor MRI approach for skeletal muscle fascicle length measurements. Physiol Rep. 2016;4(24).

17. Sinha S, Sinha U, Edgerton VR. In vivo diffusion tensor imaging of the human calf muscle. J Magn Reson Imaging. 2006;24(1):182–90.

18. Qi J, Olsen NJ, Price RR, Winston JA, Park JH. Diffusion-weighted imaging of inflammatory myopathies: polymyositis and dermatomyositis. J Magn Reson Imaging. 2008;27(1):212–7.

19. Sinha U, Yao L. In vivo diffusion tensor imaging of human calf muscle. J Magn Reson Imaging. 2002;15(1):87–95.

20. Van Donkelaar CC, Kretzers LJ, Bovendeerd PH, Lataster LM, Nicolay K, Janssen JD, Drost MR. Diffusion tensor imaging in biomechanical studies of skeletal muscle function. J Anat. 1999;194 ( Pt 1)(Pt 1):79–88.

21. Guttsches AK, Rehmann R, Schreiner A, Rohm M, Forsting J, Froeling M, et al. Quantitative Muscle-MRI Correlates with Histopathology in Skeletal Muscle Biopsies. J Neuromuscul Dis. 2021;8(4):669–78.

22. Berry DB, Englund EK, Galinsky V, Frank LR, Ward SR. Varying diffusion time to discriminate between simulated skeletal muscle injury models using stimulated echo diffusion tensor imaging. Magn Reson Med. 2021;85(5):2524–36.

23. Sigmund EE, Novikov DS, Sui D, Ukpebor O, Baete S, Babb JS, et al. Time-dependent diffusion in skeletal muscle with the random permeable barrier model (RPBM): application to normal controls and chronic exertional compartment syndrome patients. NMR Biomed. 2014;27(5):519–28.

24. Mazzoli V, Oudeman J, Nicolay K, Maas M, Verdonschot N, Sprengers AM, et al. Assessment of passive muscle elongation using Diffusion Tensor MRI: Correlation between fiber length and diffusion coefficients. NMR Biomed. 2016;29(12):1813–24.

25. Yucesoy CA, Koopman BH, Baan GC, Grootenboer HJ, Huijing PA. Extramuscular myofascial force transmission: experiments and finite element modeling. Arch Physiol Biochem. 2003;111(4):377–88.

26. Charles JP, Moon CH, Anderst WJ. Determining Subject-Specific Lower-Limb Muscle Architecture Data for Musculoskeletal Models Using Diffusion Tensor Imaging. J Biomech Eng. 2019;141(6).

27. Arnold EM, Ward SR, Lieber RL, Delp SL. A model of the lower limb for analysis of human movement. Ann Biomed Eng. 2010;38(2):269–79.

28. Rajagopal A, Dembia CL, DeMers MS, Delp DD, Hicks JL, Delp SL. Full-Body Musculoskeletal Model for Muscle-Driven Simulation of Human Gait. IEEE Trans Biomed Eng. 2016;63(10):2068–79.

29. Delp SL, Anderson FC, Arnold AS, Loan P, Habib A, John CT, et al. OpenSim: open-source software to create and analyze dynamic simulations of movement. IEEE Trans Biomed Eng. 2007;54(11):1940–50.

30. Chen Z, Franklin DW. Musculotendon Parameters in Lower Limb Models: Simplifications, Uncertainties, and Muscle Force Estimation Sensitivity. Ann Biomed Eng. 2023;51(6):1147–64.

31. Okamoto Y, Kunimatsu A, Kono T, Nasu K, Sonobe J, Minami M. Changes in MR diffusion properties during active muscle contraction in the calf. Magn Reson Med Sci. 2010;9(1):1–8.

32. Damon BM, Froeling M, Buck AK, Oudeman J, Ding Z, Nederveen AJ, et al. Skeletal muscle diffusion tensor-MRI fiber tracking: rationale, data acquisition and analysis methods, applications and future directions. NMR Biomed. 2017;30(3).

33. Mazzoli V, Moulin K, Kogan F, Hargreaves BA, Gold GE. Diffusion Tensor Imaging of Skeletal Muscle Contraction Using Oscillating Gradient Spin Echo. Front Neurol. 2021;12:608549.

34. Deffieux T, Gennisson JL, Tanter M, Fink M. Assessment of the mechanical properties of the musculoskeletal system using 2-D and 3-D very high frame rate ultrasound. IEEE Trans Ultrason Ferroelectr Freq Control. 2008;55(10):2177–90.

35. Gijsbertse K, Goselink R, Lassche S, Nillesen M, Sprengers A, Verdonschot N, et al. Ultrasound Imaging of Muscle Contraction of the Tibialis Anterior in Patients with Facioscapulohumeral Dystrophy. Ultrasound Med Biol. 2017;43(11):2537–45.

36. Gijsbertse K, Sprengers AM, Nillesen MM, Hansen HH, Lopata RG, Verdonschot N, de Korte CL. Three-dimensional ultrasound strain imaging of skeletal muscles. Phys Med Biol. 2017;62(2):596–611.

37. Raiteri BJ, Cresswell AG, Lichtwark GA. Three-dimensional geometrical changes of the human tibialis anterior muscle and its central aponeurosis measured with three-dimensional ultrasound during isometric contractions. PeerJ. 2016;4:e2260.

38. Yaman A, Ozturk C, Huijing PA, Yucesoy CA. Magnetic resonance imaging assessment of mechanical interactions between human lower leg muscles in vivo. J Biomech Eng. 2013;135(9):91003.

39. Pamuk U, Karakuzu A, Ozturk C, Acar B, Yucesoy CA. Combined magnetic resonance and diffusion tensor imaging analyses provide a powerful tool for in vivo assessment of deformation along human muscle fibers. J Mech Behav Biomed Mater. 2016;63:207–19.

40. Williams SE, Heemskerk AM, Welch EB, Li K, Damon BM, Park JH. Quantitative effects of inclusion of fat on muscle diffusion tensor MRI measurements. J Magn Reson Imaging. 2013;38(5):1292–7.

41. Damon BM, Ding Z, Hooijmans MT, Anderson AW, Zhou X, Coolbaugh CL, et al. A MATLAB toolbox for muscle diffusion-tensor MRI tractography. J Biomech. 2021;124:110540.

42. Yushkevich PA, Piven J, Hazlett HC, Smith RG, Ho S, Gee JC, Gerig G. User-guided 3D active contour segmentation of anatomical structures: significantly improved efficiency and reliability. Neuroimage. 2006;31(3):1116–28.

43. Thirion JP. Image matching as a diffusion process: an analogy with Maxwell’s demons. Med Image Anal. 1998;2(3):243–60.

44. Vercauteren T, Pennec X, Perchant A, Ayache N. Diffeomorphic demons: efficient non-parametric image registration. Neuroimage. 2009;45(1 Suppl):S61–72.

45. Xu Q, Anderson AW, Gore JC, Ding Z. Efficient anisotropic filtering of diffusion tensor images. Magn Reson Imaging. 2010;28(2):200–11.

46. Ding Z, Gore JC, Anderson AW. Reduction of noise in diffusion tensor images using anisotropic smoothing. Magn Reson Med. 2005;53(2):485–90.

47. Buck AK, Ding Z, Elder CP, Towse TF, Damon BM. Anisotropic Smoothing Improves DT-MRI-Based Muscle Fiber Tractography. PLoS One. 2015;10(5):e0126953.

48. Damon BM, Heemskerk AM, Ding Z. Polynomial fitting of DT-MRI fiber tracts allows accurate estimation of muscle architectural parameters. Magn Reson Imaging. 2012;30(5):589–600.

49. Lockard C. Impact of tract propogation stop criteria on skeletal muscle diffusion-tensor-imaging fiber completeness and characteristics. In: Hooijmans MT, editor. Proceedings International Society of Magnetic Resonance Imaging 2022.

50. Sacks RD, Roy RR. Architecture of the hind limb muscles of cats: functional significance. J Morphol. 1982;173(2):185–95.

51. Ding Z. Case study: reconstruction, visualization and quantification of neuronal fiber pathways. In: Gore JC, editor. Proceedings Visualization: VIS’01., 2001•ieeexplore.ieee.org; 2001.

52. Heemskerk AM, Damon BM. Diffusion Tensor MRI Assessment of Skeletal Muscle Architecture. Curr Med Imaging Rev. 2007;3(3):152–60.

53. Bolsterlee B, D’Souza A, Herbert RD. Reliability and robustness of muscle architecture measurements obtained using diffusion tensor imaging with anatomically constrained tractography. J Biomech. 2019;86:71–8.

54. Charles JP, Suntaxi F, Anderst WJ. In vivo human lower limb muscle architecture dataset obtained using diffusion tensor imaging. PLoS One. 2019;14(10):e0223531.

55. Maganaris CN, Baltzopoulos V. Predictability of in vivo changes in pennation angle of human tibialis anterior muscle from rest to maximum isometric dorsiflexion. Eur J Appl Physiol Occup Physiol. 1999;79(3):294–7.

56. Chleboun GS, Busic AB, Graham KK, Stuckey HA. Fascicle length change of the human tibialis anterior and vastus lateralis during walking. J Orthop Sports Phys Ther. 2007;37(7):372–9.

57. Klimstra M, Dowling J, Durkin JL, MacDonald M. The effect of ultrasound probe orientation on muscle architecture measurement. J Electromyogr Kinesiol. 2007;17(4):504–14.

58. de Boer MD, Seynnes OR, di Prampero PE, Pisot R, Mekjavic IB, Biolo G, Narici MV. Effect of 5 weeks horizontal bed rest on human muscle thickness and architecture of weight bearing and non-weight bearing muscles. Eur J Appl Physiol. 2008;104(2):401–7.

59. Ward SR, Eng CM, Smallwood LH, Lieber RL. Are current measurements of lower extremity muscle architecture accurate? Clin Orthop Relat Res. 2009;467(4):1074–82.

60. Wickiewicz TL, Roy RR, Powell PL, Edgerton VR. Muscle architecture of the human lower limb. Clin Orthop Relat Res. 1983(179):275–83.

61. Lieber RL, Friden J. Clinical significance of skeletal muscle architecture. Clin Orthop Relat Res. 2001(383):140–51.

62. Powell PL, Roy RR, Kanim P, Bello MA, Edgerton VR. Predictability of skeletal muscle tension from architectural determinations in guinea pig hindlimbs. J Appl Physiol Respir Environ Exerc Physiol. 1984;57(6):1715–21.

63. Secondulfo L, Hooijmans MT, Suskens JJ, Mazzoli V, Maas M, Tol JL, et al. A diffusion tensor-based method facilitating volumetric assessment of fiber orientations in skeletal muscle. PLoS One. 2022;17(1):e0261777.

64. Deux JF, Malzy P, Paragios N, Bassez G, Luciani A, Zerbib P, et al. Assessment of calf muscle contraction by diffusion tensor imaging. Eur Radiol. 2008;18(10):2303–10.

65. Oudeman J, Nederveen AJ, Strijkers GJ, Maas M, Luijten PR, Froeling M. Techniques and applications of skeletal muscle diffusion tensor imaging: A review. J Magn Reson Imaging. 2016;43(4):773–88.

66. Huijing PA, Yaman A, Ozturk C, Yucesoy CA. Effects of knee joint angle on global and local strains within human triceps surae muscle: MRI analysis indicating in vivo myofascial force transmission between synergistic muscles. Surg Radiol Anat. 2011;33(10):869–79.

67. Wang H, Dong L, O’Daniel J, Mohan R, Garden AS, Ang KK, et al. Validation of an accelerated ‘demons’ algorithm for deformable image registration in radiation therapy. Phys Med Biol. 2005;50(12):2887–905.

68. Latifi K, Zhang G, Stawicki M, van Elmpt W, Dekker A, Forster K. Validation of three deformable image registration algorithms for the thorax. J Appl Clin Med Phys. 2013;14(1):3834.

69. Zhou X. Predicted effects of Image Acquisition and Analysis Conditions on muscle DTMRI Tractography-Based Architectural Estimates. In: Lockard C, editor. Proceedings International Society of Magnetic Resonance Imaging in Medicine 2023.

70. Damon BM. Effects of image noise in muscle diffusion tensor (DT)-MRI assessed using numerical simulations. Magn Reson Med. 2008;60(4):934–44.

71. Froeling M, Nederveen AJ, Nicolay K, Strijkers GJ. DTI of human skeletal muscle: the effects of diffusion encoding parameters, signal-to-noise ratio and T2 on tensor indices and fiber tracts. NMR Biomed. 2013;26(11):1339–52.

